# FUS binding to RNA prevents R-loops

**DOI:** 10.1101/2022.08.11.503633

**Authors:** Valery F. Thompson, Daniel R. Wieland, Vivian Mendoza-Leon, Helen I. Janis, Michelle A. Lay, Lucas M. Harrell, Jacob C. Schwartz

## Abstract

The protein FUS (FUSed in sarcoma) is a metazoan RNA-binding protein that influences RNA production by all three nuclear polymerases. FUS also binds nascent transcripts, RNA processing factors, RNA polymerases, and transcription machinery. We explored the role of FUS binding interactions for activity during transcription. *In vitro* run-off transcription assays revealed FUS enhanced RNA produced by a non-eukaryote polymerase. Activity also reduced the formation of R-loops between RNA products and their DNA template. Analysis by domain mutation and deletion indicated RNA-binding was required for activity. We interpret that FUS binds and sequesters nascent transcripts to prevent R-loops forming with nearby DNA. DRIP-seq analysis showed that a knockdown of FUS increased R-loop enrichment near expressed genes. Prevention of R-loops by FUS binding to nascent transcripts has potential to affect transcription by any RNA polymerase, highlighting the broad impact FUS can have on RNA metabolism in cells and disease.

## INTRODUCTION

A prominent member of the heterogeneous nuclear ribonucleoprotein (hnRNP) family is FUS (FUsed in Sarcoma). FUS is best known for gene mutations leading to amyotrophic lateral sclerosis or pediatric sarcomas (1–3). FUS is conserved throughout metazoan species and among the highest expressed proteins in human tissues (3). FUS has a structure that is largely intrinsically disordered (4). The disordered domains contained in FUS are its low complexity (LC) domain and three arginine and glycine-rich (RGG/RG) domains. Evidence of FUS activity on transcription is reported for all three RNA polymerases in the nucleus (5–9). FUS binds directly to nascent RNA transcripts, RNA Pol II, RNA Pol III, and other transcription and RNA-processing factors (4,10–13). Outside of these, FUS also interacts with mRNA, DNA repair machinery, DNA loops formed during recombination, and snoRNAs (12,14–16).

FUS affinity for RNA involves an RNA-recognition motif (RRM), Zinc finger domain (ZnF), and the three disordered RGG/RG domains (17–19). FUS binds a large number and variety of messenger and noncoding RNAs with a degenerate specificity but prefers G-rich sequences, RNA stem-loops, and more complex RNA structures (18–20). RNA binding drives FUS to oligomerize into large assemblies, which enhance binding to RNA Pol II and other protein partners (21,22). In cells, the assembly behavior of FUS is found in protein particles that contain FUS and RNA Pol II and form in a transcription-dependent manner (5,23). FUS is also enriched near transcription start sites (TSS) of most expressed genes encoding mRNA and prevents phosphorylation of Ser-2 (Ser2P) in the heptad repeat of the C-terminal domain (CTD) in RNA Pol II (8,24).

We set out to better define the role that binding the RNA polymerase has in FUS activity on transcription. We anticipated direct binding to the polymerase and RNA processing factors would be required for FUS activity (7,8,17). However, FUS showed activity on the transcription of a non-eukaryotic RNA polymerase intended to be a negative control. We decided to refocus our study on this unexpected activity observed in the absence of previously known protein:protein interactions.

## MATERIAL AND METHODS

### Recombinant protein expression and purification

FUS constructs used in this study are available upon request. All recombinant proteins were expressed and purified from *E. coli* BL21(DE3) cells. All FUS constructs were N-terminally fused to 6x His and maltose-binding protein (MBP) to improve purification and solubility. Protein expression was induced after cell growth reached an OD600 of 0.8 and then allowed to continue overnight at 17 °C with shaking (200 rpm). Purification was made using 1 to 2 g of frozen pellets of induced *E. coli*, lysates incubated with Ni-NTA Sepharose beads (Cytiva Lifesciences, 17531802), and eluted in FUS-SEC buffer (1 M urea, 0.25 M KCl, 50 mM Tris-HCl pH 8.0) with 250 mM imidazole and either 1.5 mM β-mercaptoethanol or 1 mM DTT added (5,18,20).

Plasmids for protein expression of T7(P266L) polymerase fused with a 6xHis tag were provided by A. Berman (University of Pittsburgh) (25). Expression in *E. coli* BL21(DE3) cells and purification were done essentially as previously published (26). Protein expression was induced by 0.5 mM Isopropyl β-D-1-thiogalactopyranoside (IPTG) when transformed cells grew to an OD600 of 0.6 then continued for 3 hours at 37 °C. Induced cells stored at −80 °C were thawed, lysed in T7 Buffer (50 mM Tris-HCl pH 8.0, 100 mM NaCl, 5% v/v glycerol, and 5 mM β-mercaptoethanol added just before use) with 1 mM imidazole then incubated with Ni-NTA beads. T7 Buffer with increasing imidazole was used to wash beads 4 times (1 mM imidazole), 2 times (10 mM imidazole), and to elute protein (100 mM imidazole).

Size-exclusion chromatography (SEC) was performed with a 10/300 Superdex® 200 column (Cytiva Lifesciences, 17517501) for FUS using FUS-SEC buffer and for T7 Pol with T7-SEC buffer (20 mM Tris-HCl pH 7.5, 100 mM NaCl, 0.1 mM EDTA, and 1 mM DTT). FUS protein was stored at room temperature for up to 4 weeks or until activity appeared lost. T7 Pol activity was assessed by titrating protein into transcription assays as described below. The absence of nuclease activity was confirmed by comparing plasmid DNA and transcribed RNA incubated overnight with or without protein and electrophoresis using a Novex™ TBE-urea, 6% polyacrylamide gel (Invitrogen, EC68655BOX) and stained with SYBR-Gold (Invitrogen, S11494).

### Nucleic Acid preparation and purification

DNA substrates used in T7 transcription assays were prepared from pcDNA3 vector plasmid and gene inserted for RNA transcription was Flag-FUS (18). Bsu36I restriction enzyme (50 U, NEB, R0524S) was used to linearize plasmid (10 µg) by incubating at 37 °C for a minimum of 2 hours in 1x CutSmart buffer (NEB, B7204S). Linearized plasmid was purified by extraction with an equal volume of phenol:chloroform:isoamyl alcohol (25:24:1, pH 6.7/8.0) (Fisher Scientific, BP1752I-100) followed by two additional chloroform extractions (VWR, 97064-680) and ethanol precipitated. Plasmid concentrations were measured by UV absorption (260 nm) using a Biotek Epoch 2 plate reader with a TAKE3 plate. RNA TET456 was provided by T.R. Cech (University of Colorado Boulder) (27). Additional nucleic acids were yeast tRNA (Invitrogen, 15401011) and a commercially synthesized ssDNA (Millipore-Sigma). Nucleic acid sequences can be found in **Supplemental Table 1**.

### T7 Transcription

Transcription reactions were prepared at room temperature in 20 µL transcription buffer (40 µM Tris-HCl pH 8, 5 mM DTT, 10 mM MgCl_2_) with 2 mM NTP’s (NEB, N0450S) and 40 U/µL RNasin™ Plus (Promega, N2615). T7 (P266L) polymerase (28) was added to linearized plasmid DNA (25 ng/µL) at 37 °C, incubated for up to 2 hours, and halted either by heating to >75 °C or addition of EDTA (12 mM final concentration). To detect RNA products by electrophoresis, samples were heated (90 °C, 5 min) in formamide loading dye (49% formamide, 5 mM EDTA, 0.05% xylene cyanol, 0.05% bromophenol blue) and loaded to a Novex™ TBE-Urea, 6% polyacrylamide gels (Invitrogen, EC68655BOX) to run in TBE (VWR, 97061-754) at 180 V for 50 minutes. A ssRNA ladder (NEB, N0364S) was used for molecular weight standards. Gels were stained with SYBR-Gold (Invitrogen, S11494) in TBE and imaged using a Bio-Rad ChemiDoc™ MP Imaging System. Densitometry was performed with Bio-Rad Image Lab™ Software v. 6.0.1. The ratio of RNA/DNA was calculated for each sample to control for loading differences. RNA and DNA were also quantified by Quant-iT™ Assay (Thermo Fisher Scientific, Q333140 and Q33120) according to manufacturer’s instructions.

### Co-immunoprecipitation assay

Recombinant expressed FUS protein (1.3 µg) and anti-FUS (4H11) antibody (11 µg; Santa Cruz Biotechnology, sc-47711) were immobilized by incubating together for 1 hour at room temperature in phosphate buffered saline (PBS, pH 7.4) (Sigma, P3813) and 0.1 mg/mL bovine serum albumin protein (VWR, 97062-508), then 1 hour with Pierce™ Protein A/G agarose beads (Thermo Fisher Scientific, 20421). Beads were washed 4 times with 0.1% NP-40 substitute (VWR, 97064-730) in tris-buffered saline (TBS). Beads with FUS were incubated with T7 Pol for 1 hour at room temperature with rotation, then washed twice in high salt, twice in TBS with 0.1% NP-40 substitute, and one with TBS. Elution of protein bound to beads was incubated at 95 °C for ten minutes in 2x NuPAGE® LDS sample loading buffer (Life Technologies, NP0008) containing 4% lithium dodecyl-sulfate (LDS) and DTT (100 mM) added. Nucleic acids used were prepared as described above and **Supplemental Table 1**. Detection of eluted proteins was made by western blot assay using anti-FUS (A300-294A, Bethyl Laboratories) to identify FUS protein and anti-His (HIS.H8, Novus Biologicals, NBP2-31055) to identify His tags for both FUS and T7 Pol proteins.

### RNA:DNA hybrid dot blot assays

RNA:DNA hybrids were detected and quantified by dot blot assay. From transcription assays, 10% of reactions were spotted on N+ membranes (Fisher Scientific, 45-000-850), dried and UV crosslinked. Blots were blocked with 5% nonfat dried milk (NFDM) in TBS-T, then incubated overnight at 4 °C with an anti-RNA:DNA hybrid antibody S9.6 against RNA:DNA hybrids (1/2000 dilution) in TBS-T with 2.5% NFDM. Blots were washed 5 minutes 4 times in TBS-T, incubated in secondary antibody (goat anti-mouse IgG-HRP, 1/20000) for 1 hr at RT. Images were taken after incubation with SuperSignal™ West Pico PLUS (Fisher Pierce, PI34578) and imaged using a Chemidoc MP system (BioRad).

### DRIP-seq assay and data analysis

DRIP-seq assays were performed essentially as a previously published protocol (29). HEK293T/17 cells (ATCC, CRL-11268) grown and passaged in DMEM (5% FBS) were transfected by siRNA using RNAiMax™ (Invitrogen, 13778150) in Optimem (Invitrogen, 31985070). Sequences of siRNAs used are included in **Supplemental Table 1**. Between 7 and 9 x 10^6^ cells were harvested 72 hours post-siRNA treatment, and cell pellets were frozen in liquid nitrogen. Pellets were later resuspended in 1.6 mL of TE buffer (10 mM Tris-HCl pH 8, 1 mM EDTA) that was added to include 0.6% sodium dodecyl sulfate (SDS) and 60 µg/mL Proteinase K. After Proteinase K digested lysates overnight at 37 °C, DNA was extracted using the MaXtract High Density tubes (Qiagen, 129065). DNA was spooled by glass Pasteur pipette, transferred, then ethanol precipitated.

DNA was digested overnight in TE buffer with 1x NEBuffer™ 2 (New England Biolabs Inc., #B6002), BSA (95 µg/mL), spermidine (1 mM), and 30 U each of restriction enzymes BsrGl, EcoRI, HindIII, SspI, and XbaI. Restriction digest was confirmed by agarose (0.8% w/v) gel electrophoresis. After restriction enzyme digestions, DNA was extracted by Phase Lock Gel™ (VWR, 10847-800) per manufacturer instructions. Negative control samples were digested by 40U RNaseH (Fisher Scientific, 50-811-717) per 10 µg DNA. 50 µg of DNA was incubated overnight at 4 °C with 20 µg S9.6 antibody and in 500 µL DRIP buffer (1x TE buffer, 10 mM Na phosphate pH 7, 140 mM NaCl, 0.05% Triton X-100). 5 µg of DNA was kept for input samples. Antibody precipitated with 100 µL Protein A/G beads (Millipore, IP05-1.5ML) was washed twice in 700 µL DRIP buffer. Hybrids were eluted by incubating beads with 300 µL DRIP Elution buffer (50 mM Tris-HCl pH 8, 10 mM EDTA, 0.5% SDS, 0.5 µg/µL Proteinase K) for 45 minutes at 55 °C, then extracted by Phase Lock Gel™. Samples were sonicated for 30 cycles (30 sec on / 30 sec off) follow by ethanol precipitation.

Sample libraries were prepared and sequenced by Novogene Corporation Inc. using a NovoSeq6000 with 150 base paired-end reads (8 G raw data per sample). Reads were trimmed using TrimGalore (version 0.6.6) and aligned to GRCh38 using STAR (version_2.7.8a), essentially as previously described. Bam files were processed using the deepTools2.0 suite (version 3.5.1) to normalize to 1x depth (reads per genome coverage) with bamCoverage, merge replicates by bigwigCompare, and tags binned by computeMatrix, which was then used to compute scaling factors and generate heat maps and profiles. Merged bigwig files were converted to bedgraph and then peaks by Macs2 (version 2.1.1.20160309). HOMER (version v4.11.1) was used to calculate GC-content and annotate peaks. Data was visualized, and figures were generated using Integrated Genomics Viewer (version 2.5.0). DRIP-seq data is available (GSE206740) from the Gene Expression Omnibus (https://www.ncbi.nlm.nih.gov/geo/).

## RESULTS

### FUS increases RNA yields from T7 Pol transcription

In our previous work, we observed FUS at concentrations found in the cell, >2 μM, greatly increased RNA Pol II transcription from a naked DNA template (5). We asked whether the specific protein-protein interactions FUS makes with RNA Pol II were required for activity. We chose the T7 phage RNA polymerase (T7 Pol) as a negative control that lacked binding sites present in mammalian RNA polymerases (28). Recombinantly expressed FUS with an N-terminal maltose binding protein (MBP) tag and T7 Pol were purified to homogeneity for *in vitro* run-off transcription of a linear DNA template (**Supplemental Figure 1A**).

Accumulated products of run-off transcription were observed by polyacrylamide gel electrophoresis (PAGE) and SYBR staining (**Figure 1A**). Two stained bands could be seen: a larger band that was confirmed by DNase I digestion to be linear DNA template (**Supplemental Figure 1B–C**), and a smaller band confirmed by RNaseA digestion to be the RNA transcript (**Supplemental Figure 1D**). The DNA template detected offered a normalization control for quantitative analysis of PAGE images.

**Figure 1.**
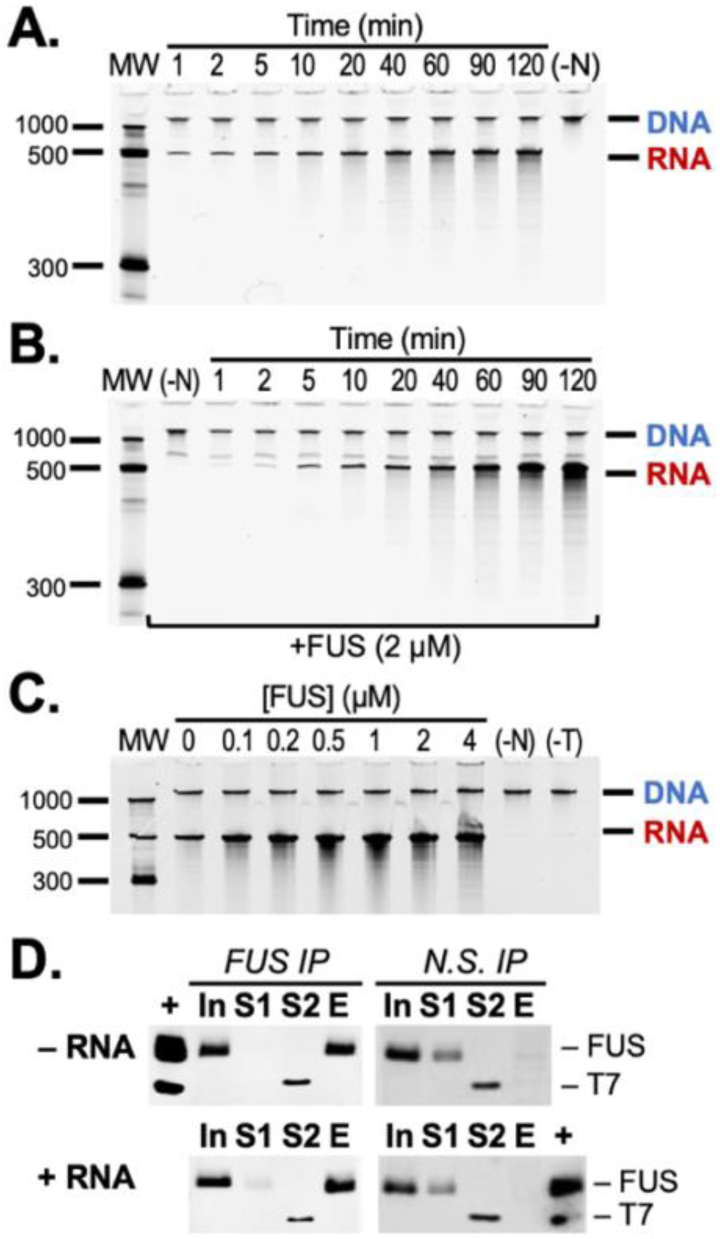
FUS increases RNA yield of run-off transcription by T7 Pol. (A) RNA product of T7 Pol run-off transcription seen by urea-PAGE and SYBR staining. The upper bands show the DNA template transcribed. Lower bands show the increase in RNA product over time. “MW” indicates molecular weight ladder. “(-N)” indicates a negative control without nucleotide triphosphates present. **(B)** RNA product of T7 Pol transcription is greater in the presence of FUS (2 µM). **(C)** RNA product is increased as a function of FUS concentration. The negative control “(-T)” contains FUS (4 µM), but T7 Pol is not present in the reaction. **(D)** Western assays measure FUS and T7 Pol during co-IP with nonspecific IgG or anti-FUS (4H11) antibody. Antibodies for western analysis target FUS (A300-294A) and the 6xHis tag contained in FUS and T7 Pol. Samples shown include input (In), supernatants S1 and S2 after incubation with FUS antibody and T7 Pol respectively, and elution (E) from agarose beads. Assays were performed with RNA present (+RNA) or absent (–RNA). “(+)” lanes contain FUS and T7 Pol for a positive control and molecular weight comparison. See also **Supplemental Figure 1**.

Contrary to our expectation, T7 Pol produced more RNA transcript in the presence of FUS (2 µM) (**Figure 1B**, **Supplemental Figure 1D-E**). Titrating buffer constituents present in the assay eliminated these as potential sources of activity (**Supplemental Figure 1F**). Increasing concentrations of FUS during T7 Pol transcription increased RNA products observed by PAGE (**Figure 1C**, **Supplemental Figure 1G**). Sufficient concentration of FUS produced a shift of RNA to the well, which was reversed by proteinase K treatment to eliminate FUS aggregates formed (**Supplemental Figure 1H**). FUS tendency to aggregate has also been noted by previous studies (21,30).

We investigated whether a direct interaction of FUS and T7 Pol could be observed through a co-immunoprecipitation (co-IP). The protocol used for co-IP was successful in detecting FUS binding to protein interactors, including RNA Pol II and III, in previous studies (5,9,11,30,31). T7 Pol incubated with immobilized FUS remained in the supernatant fraction and undetectable in the elution for negative controls and all replicates but one (N = 4, **Figure 1D**, **Supplemental Figure 1I**). Robust binding of T7 Pol could not be provoked by the linear DNA template, or an RNA known to cause FUS to bind RNA Pol II (**Figure 1D**, **Supplemental Figure 1I**) (8,21). We concluded that a protein:protein interaction, like that seen for FUS and RNA Pol II, was unlikely to offer a compelling mechanism for FUS to influence transcription by T7 Pol.

### Requirements for FUS activity on T7 Pol transcription

We next considered whether high protein concentrations affected transcription by non-specific molecular crowding. Increases to T7 Pol activity by molecular crowding with glycerol or PEG has been observed by previous studies (32–34). Activities were compared FUS or a well-folded soluble protein, bovine serum albumin (BSA), present at equal concentration during run-off transcription. BSA had no effect on RNA products observed (**Figure 2A**). Fluorescence-based quantification with an RNA-specific dye also detected no activity from BSA (**Figure 2B**). We also noted FUS activity was diminished after storage at −80 °C, but stored at room temperature, FUS activity remained stable up to 4 weeks, then no activity was seen at 12 weeks (**Supplemental Figure 2A**). The lack of effect on transcription by inactive FUS added support to the conclusion that the activity observed was specific.

**Figure 2.**
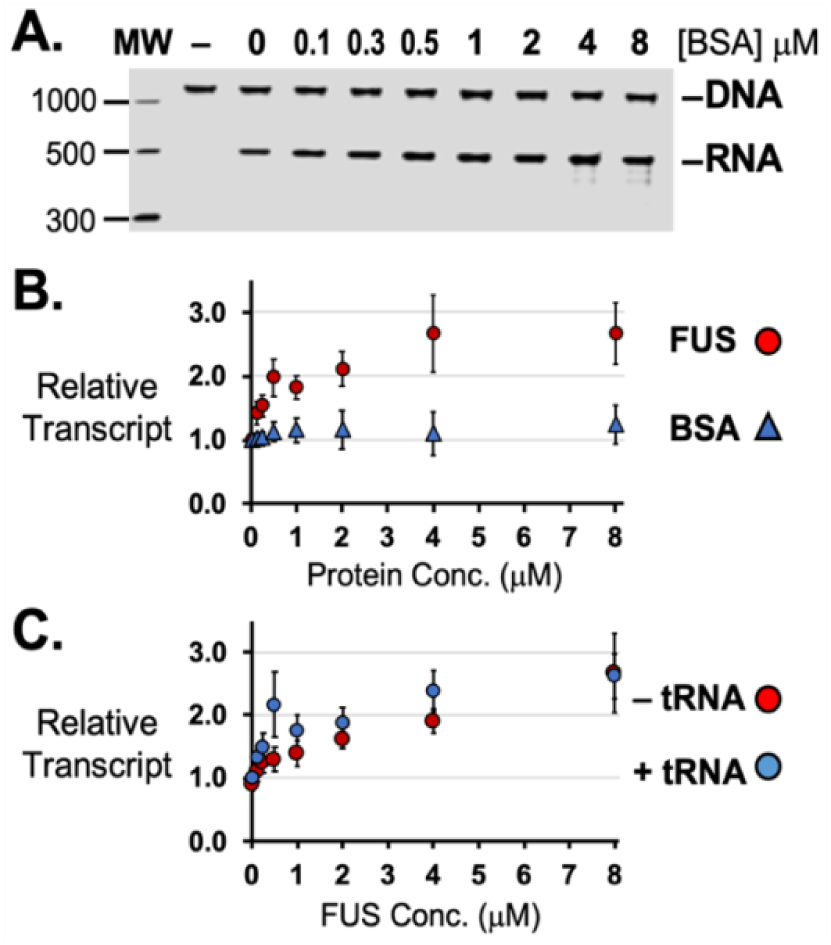
FUS effects on T7 transcription are specific. (**A**) RNA produced by T7 polymerase was not affected by titrating amounts of BSA protein. The negative control samples (–), NTPs were absent in the assay. (**B**) RNA quantified by fluorescence also show the presence or lack of increased RNA product caused by FUS or BSA, respectively. (**C**) FUS (0.1 to 8 µM) activity was unaffected with yeast tRNA (6.4 µM) present and quantified by SYBR-stained PAGE analysis to specifically measure RNA produced by run-off transcription. Results are an average of 3 or 4 experiments and relative to transcription without FUS or BSA (see also **Supplemental Figure 2**). Error bars represent ±SEM.

We considered if RNA in solution might alter FUS activity. Such an effect by RNA or DNA in solution could make the activity of FUS observed during run-off transcription difficult or impossible to observe in a cell, where concentrations of nucleic acids are high. We investigated two RNA molecules, a non-specific competitor RNA, yeast tRNA, and an RNA, TET456, previously used to induce FUS binding to RNA Pol II (21). FUS activity was unaffected by tRNA at the same concentration of transcript the assay produced (**Figure 2C**). Titrating tRNA or TET456 RNA concentrations had no effect on T7 Pol or FUS activity (**Supplemental Figure 2B-D**). A repeat of these tests with single-stranded DNA also found the activity of both proteins unaffected (**Supplemental Figure 2E**).

### FUS prevents RNA:DNA hybrids formed during transcription

Because FUS is an RNA-binding protein, we considered a mechanism for nascent RNA to affect transcription by binding its complementary template DNA to form an RNA:DNA hybrid. Together with the displaced non-template DNA strand, the structure is termed an R-loop (35,36). R-loops are found in cells at low but consistent levels, even though they can pose a threat to DNA stability, replication, and transcription (9,35,37,38). For reasons like this, cells possess multiple mechanisms to prevent or remove them (35,39).

By dot blot assay using the S9.6 antibody, RNA:DNA hybrids were found to arise early during run-off transcription and their level increased over time (**Figure 3A**) (29,40). A nuclease that cleaves RNA:DNA hybrids, RNaseH, could eliminate signals detected, confirming specificity for the S9.6 antibody in this assay. Additional controls showed no signal was produced by RNA alone or when NTP or T7 Pol was omitted from the run-off transcription assay (**Supplemental Figures 3A**).

**Figure 3.**
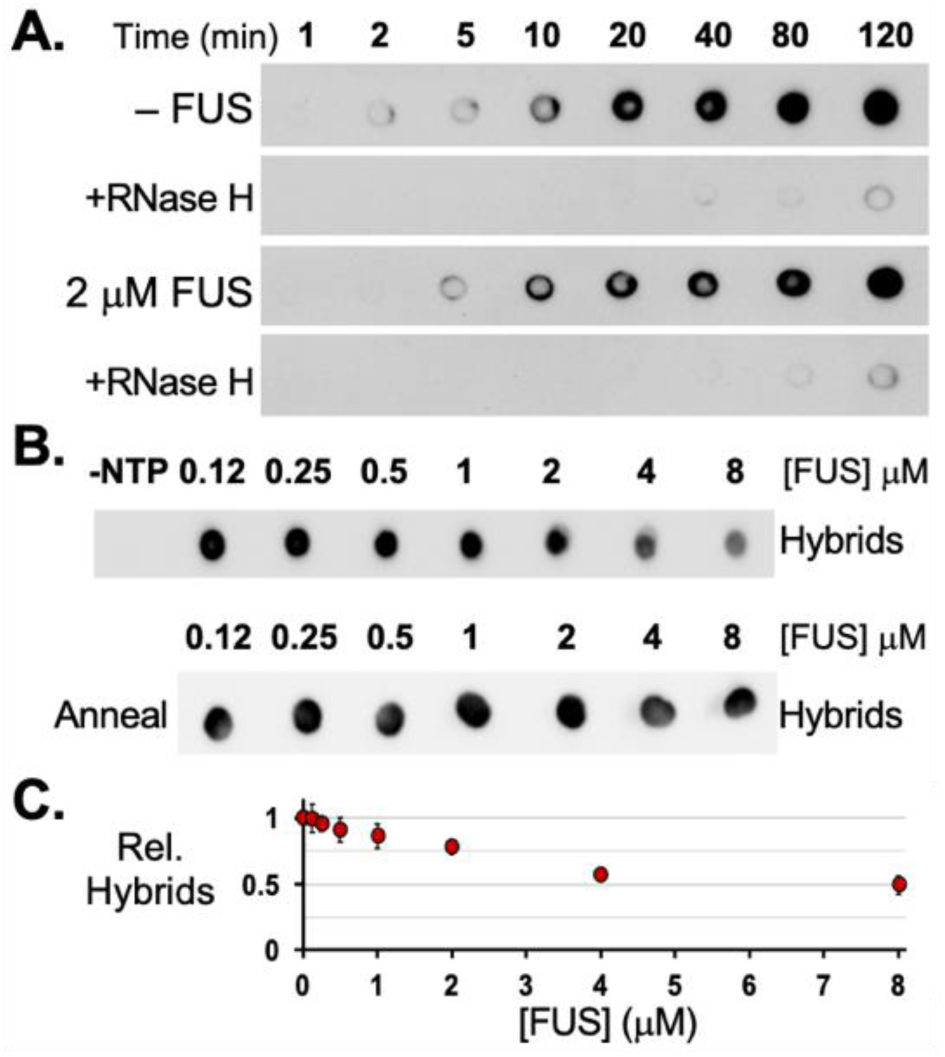
FUS prevents R-loop formation during transcription. (**A**) Dot blot assays using the S9.6 antibody detected RNA:DNA hybrids formed during T7 Pol transcription with or without FUS present (2 µM). Incubation of samples with RNaseH (+RNaseH) were used to test antibody specificity. (**B**) Dot blot assays show RNA:DNA hybrids formed at 20 minutes with increasing concentrations of FUS protein. The negative control (–NTP) has NTP omitted from the assay. As a positive control, samples were annealed by heating (95 °C) and then cooled to form hybrids. (**C**) Dot blot assays quantified by densitometry show reduced hybrids formed with FUS present relative to without FUS. Error bars represent ±SEM for 3 experiments.

The addition of FUS (2 µM) reduced RNA:DNA hybrids observed with the largest difference at 20 to 40 minutes in the assay (**Figure 3A**, **Supplemental Figure 3B**). Increasing concentration of FUS resulted in greater reduction in RNA:DNA hybrids (**Figure 3B**). Samples were heated then cooled to reveal that RNA transcribed in the presence of FUS retained their full capacity to form an RNA:DNA hybrid (**Figure 3B**, anneal). Dot blot assays could detect up to a 2-fold reduction in RNA:DNA hybrids resulted from the presence of FUS (**Figure 3C**, **Supplemental Figure 4A**).

### Low complexity and RNA-binding domains of FUS contribute to activity

We sought to identify FUS domains required for activity on T7 Pol transcription. We investigated truncations of FUS protein that lacked either the LC domain (Delta-LC) or the C-terminal RNA-binding domains (FUS-LC) (**Figure 4A**). Neither FUS-LC or Delta-LC produced a significant increase in RNA transcript or reduced RNA:DNA hybrids (**Figure 4B**, **Supplemental Figure 4A–C**). The lack of activity for Delta-LC and FUS-LC indicated that both self-assembly and RNA-binding, respectively, are required for FUS activity on T7 Pol transcription.

**Figure 4.**
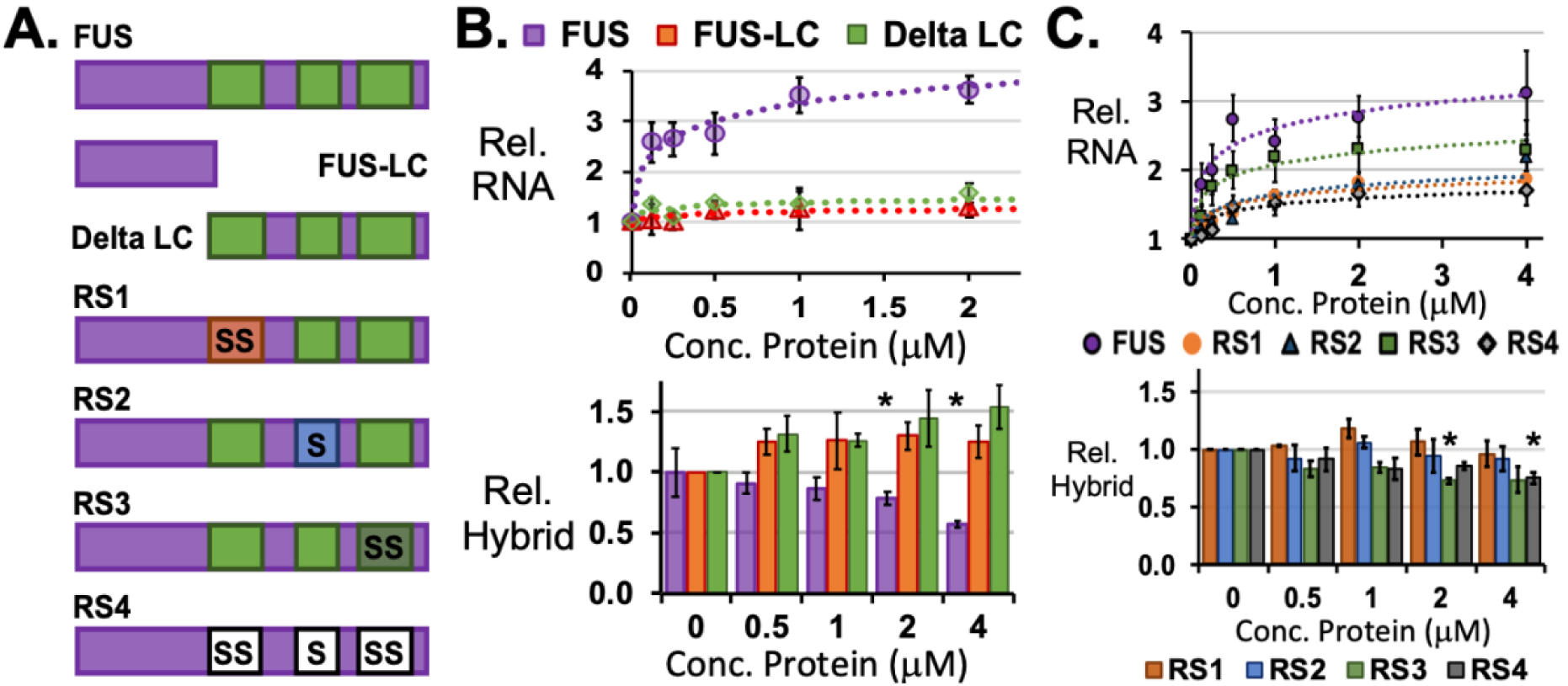
FUS oligomerization and RNA-binding domains are required to prevent R-loops. (**A**) The truncations and amino acid substitutions to FUS are illustrated in relation to its three RGG/RG domains (green). Serine residues were substituted for arginine in each RGG/RG domain (RS1, RS2, and RS3) and all RGG/RG domains (RS4). (**B**) FUS-LC and Delta-LC truncations were unable to increase RNA synthesis measured by fluorescence-based quantification or reduce RNA:DNA hybrids measured by dot blot assay. (**C**) Fluorescence-based quantification of RNA products revealed modest increases were achieved by FUS proteins with RS substitutions. The reduction in RNA:DNA hybrids seen by dot blot assays were also modest or insignificant. All results were averaged from 3 or 4 experiments. Error bars represent ±SEM. Asterisks (*) indicate p < 0.05, student t-test assuming equal variances.

We focused on the role of RNA-binding for activity by substituting serine for arginine residues in one (RS1, RS2, RS3) or all (RS4) of the RGG/RG domains in FUS (**Figure 4A**). Compared to FUS affinity for RNA, characterization of these substitutions by previous studies found RS1 and RS2 maintained similar affinity, RS3 affinity was reduced by 7-fold, and RS4 affinity was reduced by >30-fold (18). In the run-off transcription assays, RS1 or RS2 increased RNA yields up to 2-fold but did not significantly reduce RNA:DNA hybrids (**Figure 4C**, **Supplemental Figure 4D**). RS3 increased T7 Pol transcripts by >2-fold, but less than that for FUS. RNA:DNA hybrids were reduced by RS3 a small but significant amount (p < 0.05, student t-test assuming equal variances, **Figure 4C**, **Supplemental Figure 4D**). RS4 activity on transcript production matched that of RS1 and RS2, and a small reduction of hybrids could be detected (**Figure 4C**, **Supplemental Figure 4D**).

Finally, inspection by microscopy was performed to assess any self-assembled particles made by FUS or RS proteins. A necessary change from conditions in run-off transcription assays was for proteins to be allowed to incubate at room temperature for 24 hours. FUS, RS1, RS2, and RS3 proteins (2 µM) all produced visible particles up to 0.5 µm in diameter, but RS4 yielded no evidence of assemblies (**Supplemental Figure 4E**). RS4 particles were not produced by incubation of RS3 and RS4 at 10 µM (**Supplemental Figure 4F**). Lastly, FUS and RS4 were tested with protein concentrations up to 100 µM, incubation at room temperature or 4 °C, and addition of RNA, but RS4 did not yield unambiguous particles, while FUS achieved particles ≥1 µm in diameter (**Supplemental Figure 4G–H**).

### FUS expression lowers R-loop abundance in human cells

We hypothesized the activity observed for T7 Pol transcription would also be found in human cells. To test this in human HEK293T/17 cells, we used an siRNA approach to knockdown FUS expression and measured R-loop enrichment genome-wide by DRIP next-generation sequencing approach, DRIP-seq, that immunoprecipitates R-loops with the S9.6 antibody (29,41,42). Knockdown of FUS protein by siRNA (siFUS) was found to be >90% relative to control siRNA (SCR) (**Supplemental Figure 5A**) (8). Antibody detection of RNA:DNA hybrids and dsDNA in cell lysates was assessed by dot blot assay (**Supplemental Figure 5B**). Real-time PCR analysis confirmed that positive control genomic sites, *RPL13A* and *TFPT*, known to contain R-loops was enriched by S9.6 pulldown, relative to a negative control site, *EGR1*. No enrichment was observed for lysates incubated with RNaseH (**Supplemental Figure 5C**) (29). Analysis also suggested R-loop enrichment for *RPL13A* and *TFPT* increased in cells with FUS knocked down relative to control (**Supplemental Figure 5C**).

DRIP-seq analysis detected enrichment of R-loops at transcription start sites (TSS) of expressed genes, which was higher in siFUS-treated cells relative to SCR controls (**Figure 5A**). Increased enrichment of >2-fold was observed for 29% (N=5260) of protein coding genes (N=18188) (**Figure 5B**). An example typical of RNA Pol II genes, *GAPDH*, R-loop enrichment near its promoter was substantially greater in siFUS samples relative to input, SCR control, and RNaseH treated samples (**Figure 5C**) (8). The RNA Pol I genes encoding ribosomal RNA (rRNA) contained the largest increase to R-loop enrichment found in siFUS samples (**Figure 5D**).

**Figure 5.**
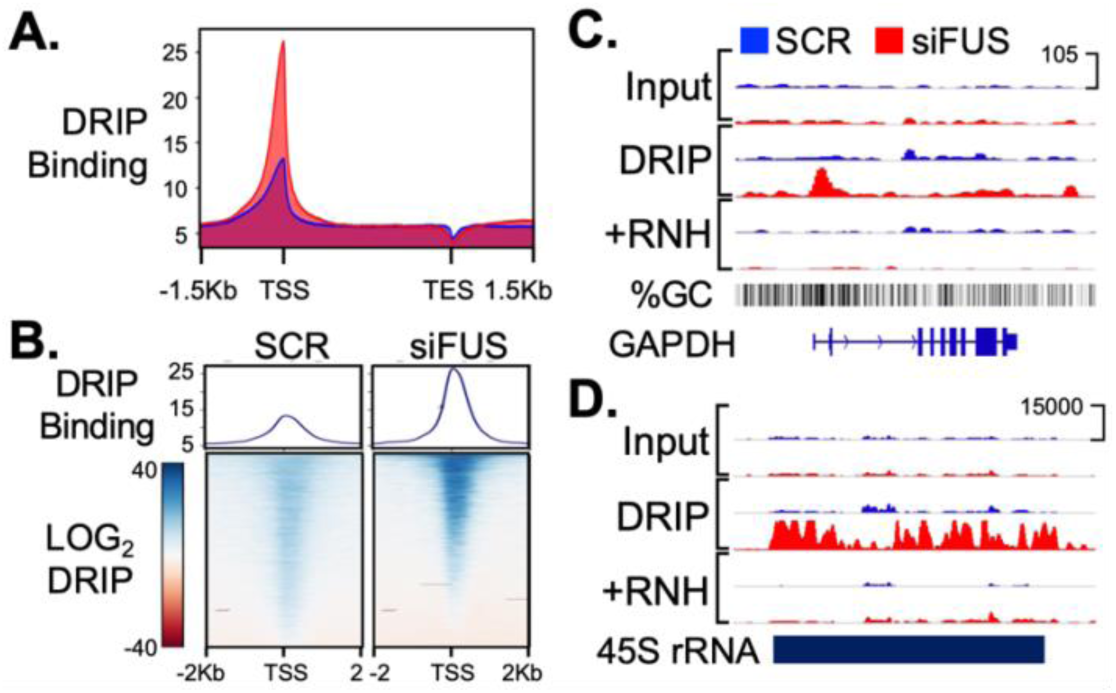
Knockdown of FUS increases R-loop abundance in human cells. DRIP-seq was performed on HEK293T/17 cells with knockdown of FUS protein, siFUS, or negative control, SCR. (**A**) Normalized DRIP-seq signal profiles at human genes were averaged and plotted for siFUS (red) or SCR (blue) samples. Gene lengths were scaled to align transcription start (TSS) and end (TES) sites. (**B**) Heat maps show DRIP-seq signals for SCR (left) or siFUS (right) treatments. (**C**) DRIP-seq showing increases to R-loop signal at the *GAPDH* gene promoter resulting from siFUS treatment. (**D**) The largest R-loop signals were found at rRNA genes in siFUS treated samples.

On average, most effects on R-loop enrichment due to FUS knockdown were in GC-rich DNA sequences (**Supplemental Figure 5D**). Also, peaks for R-loop enrichment were primarily in intergenic and satellite DNA in SCR-treated cells (N=5045 peaks), and mostly in protein-coding genes and promoters for siFUS-treated cells (N=7247 peaks) (**Supplemental Figure 5E**). Peaks produced by siFUS treatment were noted at repetitive DNA features (**Supplemental Figure 5F**). Finally, inspection of RNA Pol III transcribed genes found increased R-loop enrichment at 5S rRNA genes, but not *RMRP*, *RN7SK*, or tRNA gene clusters on chromosomes 1 and 6 (**Supplemental Figure 5G**).

In conclusion, the analysis of DRIP-seq indicated widespread FUS activity preventing R-loops in human HEK293T/17 cells. FUS activity on R-loops was not limited to transcription by RNA Pol I, II, or III. The changes to R-loop enrichment that could be measured by DRIP-seq revealed a close similarity of FUS activity in human cells and that observed during *in vitro* run-off transcription.

## DISCUSSION

Here we explore FUS activity on transcription that involves the potential for RNA transcripts to bind their DNA template, forming an R-loop. Our study followed the unexpected result that FUS increased the RNA yield from T7 Pol transcription. Subsequent analysis determined FUS reduced RNA:DNA hybrids that formed during transcription. Our model is that FUS increases RNA yield by binding nascent transcripts to prevent or delay R-loop formation, which would slow or block the RNA polymerase. In human cells, a reduction of FUS protein increased R-loop abundance. The R-loop enrichment was most prominent at expressed genes, notably those transcribed by RNA Pol I and II. These results indicate FUS bound to nascent RNA transcripts can influence transcription by preventing R-loops.

Previous studies indicate FUS influences transcription by RNA Pol I, II, and III (3,7,9,43). We have added to this list T7 Pol from bacteriophage, which shares no unambiguous sequence similarity to eukaryote RNA polymerases (44,45). Our results suggest that by preventing R-loops, FUS lowers a barrier that accumulates over the course of transcription. It is important to note that R-loops have many effects on transcription and chromatin stability besides the simple stalling of the polymerase that we have studied here (35,36,46–48). This activity for FUS also differs from those previously studied in that direct binding to the polymerase would not be a requirement for the mechanism we have proposed (21,22). The likelihood is low that a binding site is conserved between human and phage RNA polymerases, and our co-IP assay did not provide evidence of FUS binding to T7 Pol (**Figure 1D**). Binding to RNA rather than the polymerase also provides the simplest explanation for FUS to affect formation of R-loops at sites transcribed by RNA Pol I, II, and III (**Figure 5**).

Ordinarily, an increase in transcription would favor RNA:DNA hybrid formation, but the prevention of R-loops by an RNA-binding protein is not unprecedented (49). Failure to recruit snRNPs, SR proteins, and hnRNPs can allow R-loops to form (41,50–54). Protection against R-loops by SRSF1 is one example that has been reproduced in run-off transcription by T7 Pol (53). The effect seen in DRIP-seq from a FUS knockdown indicates that FUS is not easily replaced by another RNA-binding protein in the cell. This may be due in part to the contribution of the LC domain (**Figure 4B**). By comparing activity of FUS and other RNA-binding proteins with LC domains, a future study may reveal what features in a LC domain can contribute most to boost activity that prevents R-loops.

In conclusion, the ability of FUS to influence transcription can be extended to include preventing R-loops by binding nascent RNA transcripts. FUS and R-loops have complex relationships with transcription and DNA stability. The mechanism we describe would not be expected to conflict with FUS activities that involve binding the polymerase, transcription factors, or RNA-processing machinery, but this will remain to be seen by future studies. It is noteworthy that FUS and R-loops are both known to contribute to the same or related neurodegenerative diseases (9,55–58). Since FUS is highly expressed protein in human cells, its activity likely has significant implications to R-loop function and the contributions these enigmatic nucleic acid structures make to biology and disease.

## DATA AVAILABILITY

DRIP-seq data is available (GSE206740) from the Gene Expression Omnibus (https://www.ncbi.nlm.nih.gov/geo/).

## SUPPLEMENTARY DATA

Supplementary Data are available at NAR online.

## ACKNOWLEDGEMENT

We thank Dr. Andrea Berman (U. Pittsburgh) for T7 Pol plasmids and protocols; Dr. Thomas R. Cech (CU Boulder) for TET456 RNA; and H. Miller and Dr. Alex Bishop (UTHSSA) for helpful advice and discussion about DRIP-seq protocol and data analysis.

## Author contributions

V.F.T. designed and performed most experiments and analyzed data; D.R.W. performed co-IP experiments and accompanying transcription run-off and dot blot assays; V.L-M. performed transcription run-off, dot blot, and microscopy assays; H.I.J. expressed, purified, and confirmed activity of proteins; M.A.L. performed microscopy, dot blot, and western assays for DRIP-seq experiments; L.M.H. expressed, purified, and confirmed activity of proteins, performed transcription run-off assays, and optimized protocols; J.C.S. secured funding, designed and analyzed data for experiments, and supervised the project. J.C.S. wrote the manuscript. V.F.T., D.R.W., and L.M.H. edited the manuscript and writing portions of the Materials and Methods section. The final manuscript was approved by all authors.

## FUNDING

This work was supported by the National Institutes of Health [CA238499 and CA259570 to J.C.S.]; the American Cancer Society [RSG-18-237-01-DMC to J.C.S.]; and a training fellowship from the University of Arizona Cancer Center to M.A.L. Funding for open access charge: National Institutes of Health.

## CONFLICT OF INTEREST

There are no conflicts of interest to report relating to this work.

---

**Supplemental Table 1.**
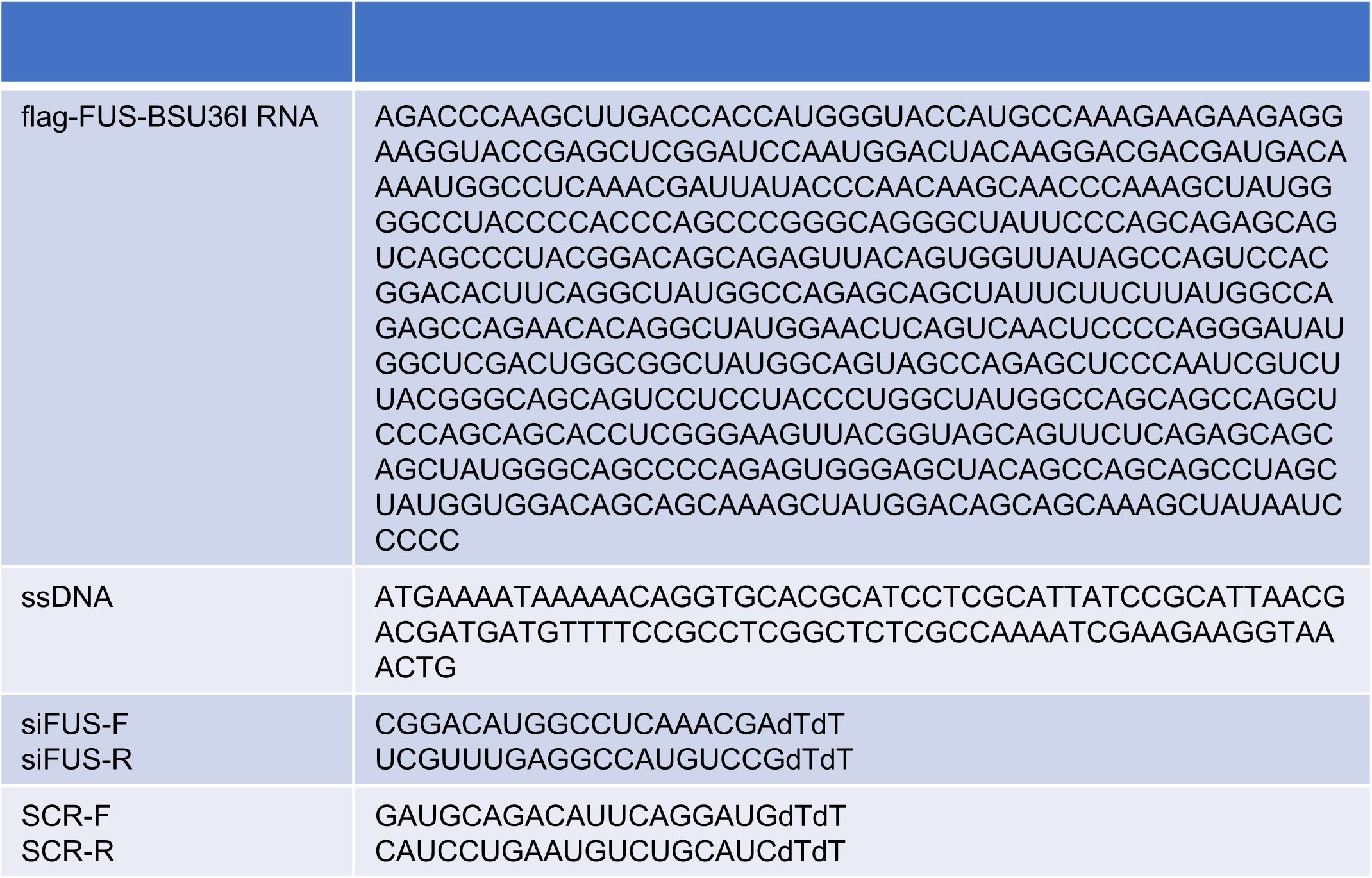

**Supplemental Figure 1.**
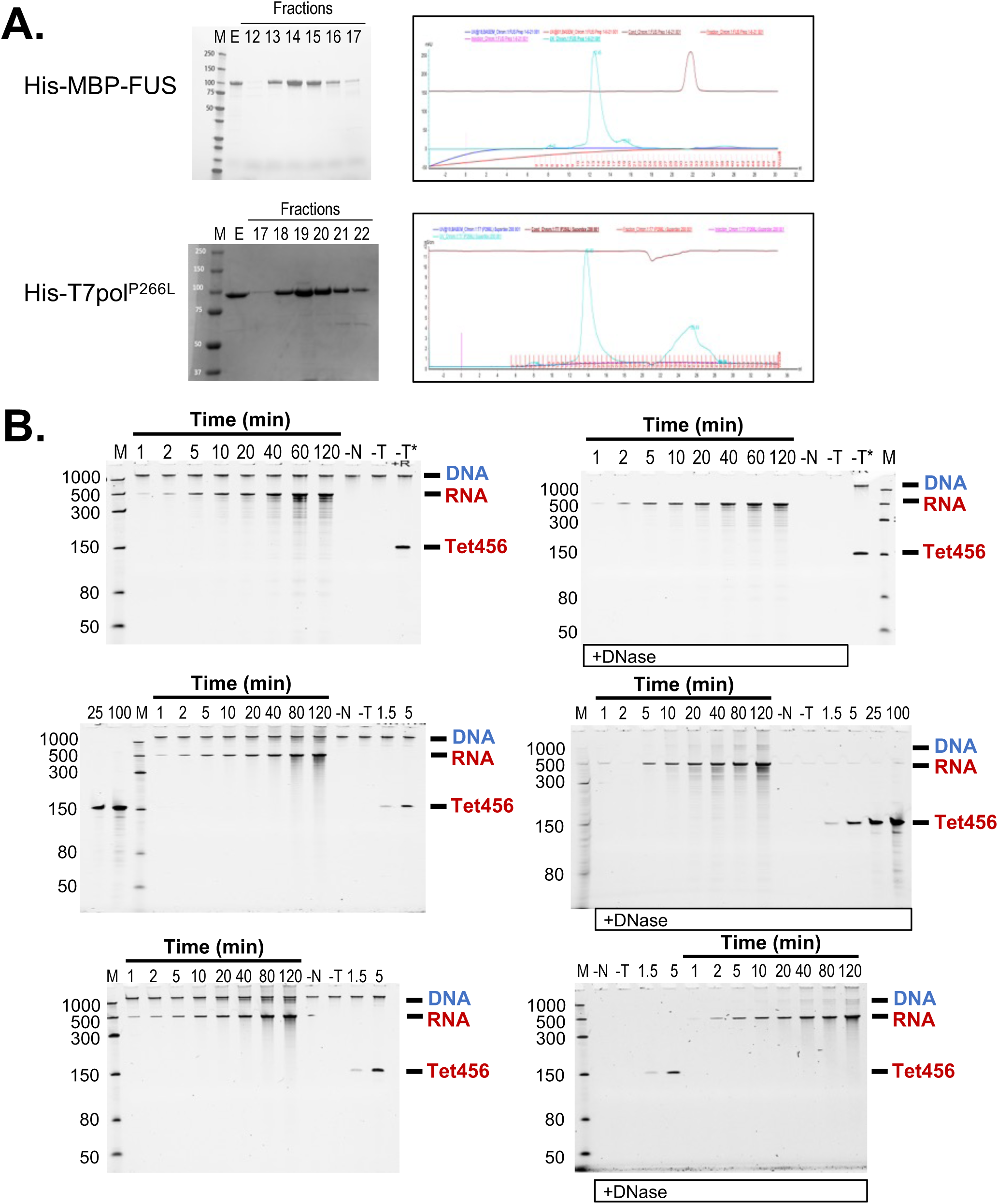
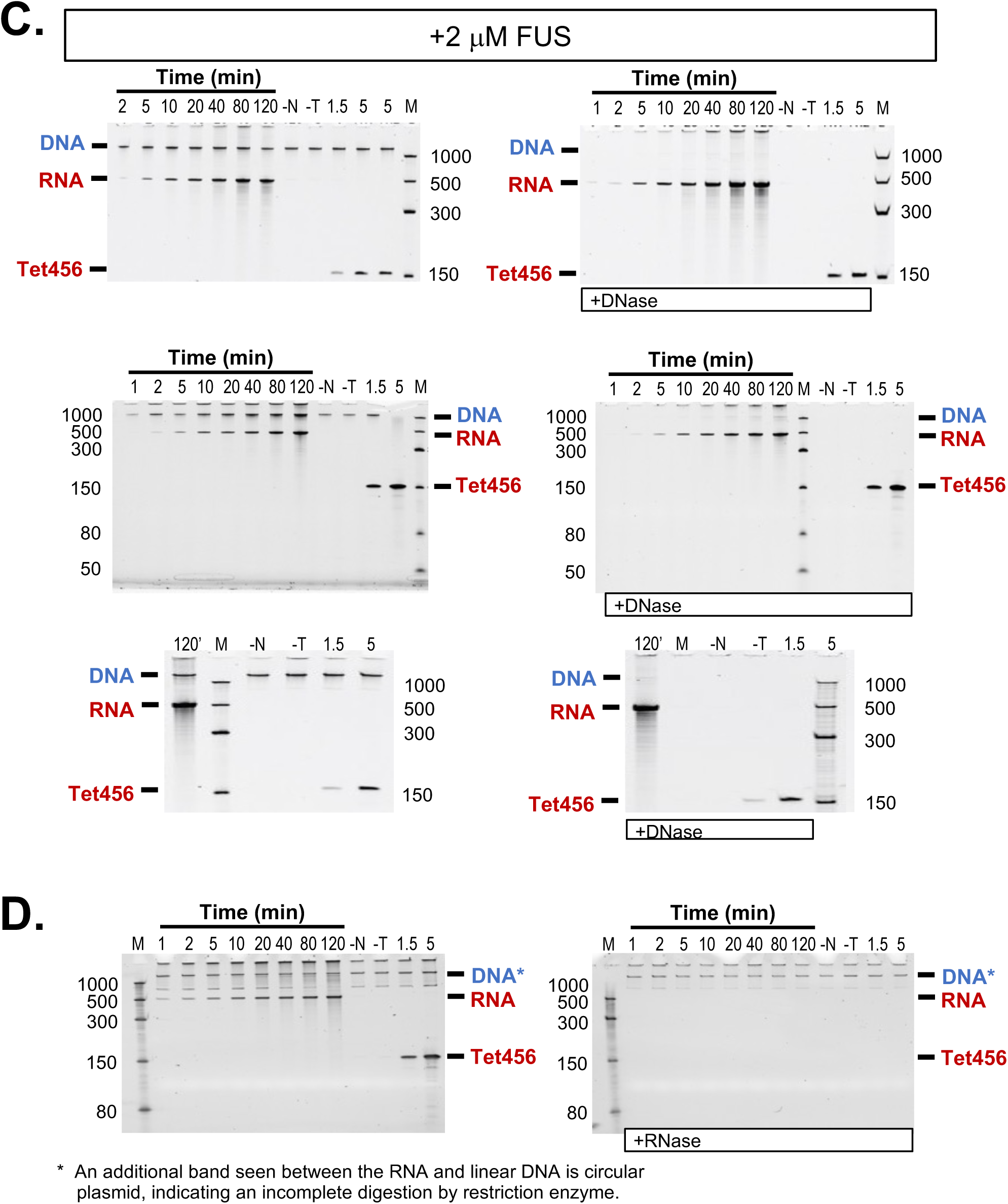
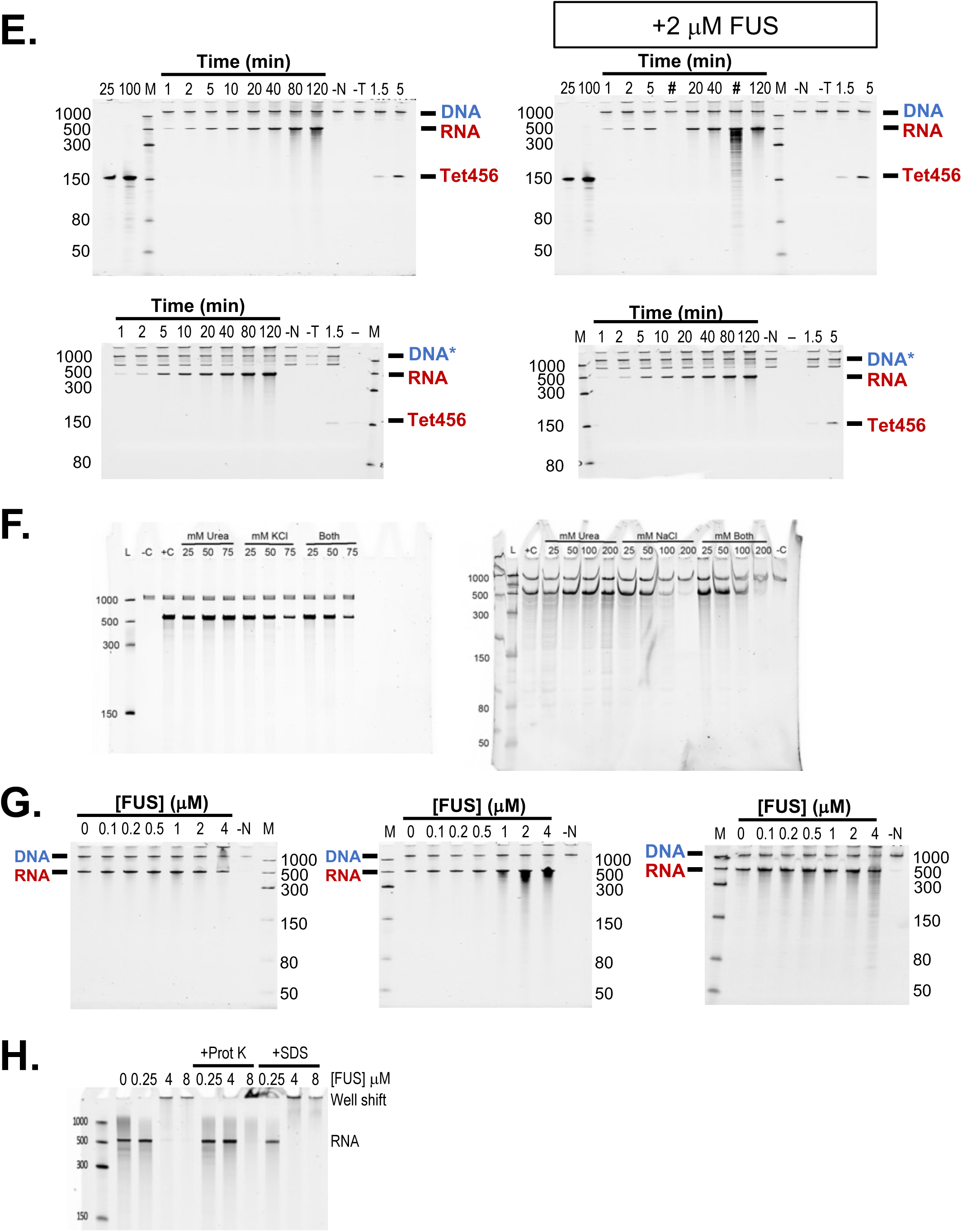
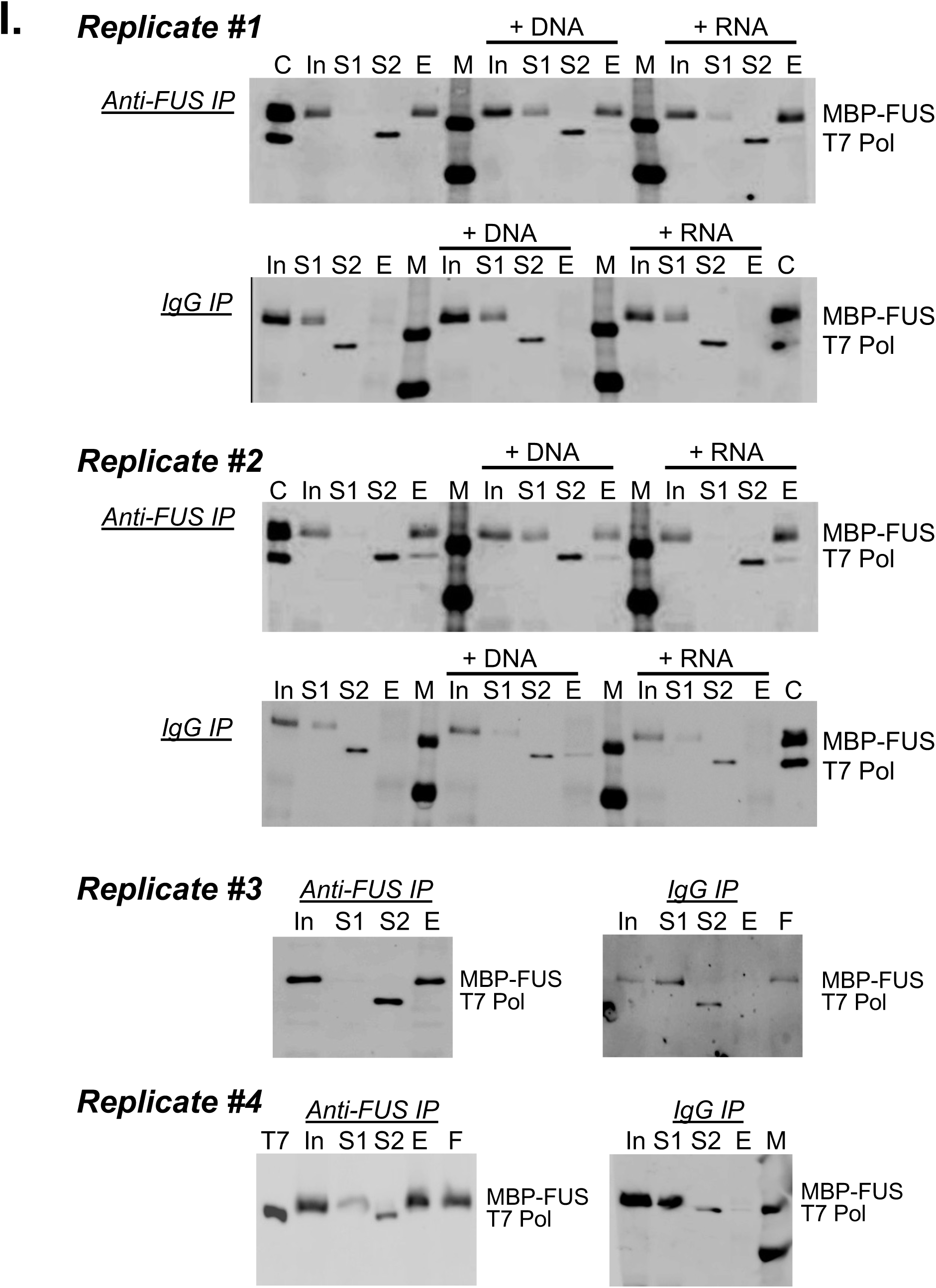
Purification of protein and immunoprecipitation. (**A**) Coomassie stained SDS-PAGE gel, left, of fractions collected after separation by SEC (see UV absorption trace, right) of His-MBP-FUS and His-T7pol^P266L^ proteins. Following SEC, FUS proteins and T7 polymerase were tested for transcription activity and presence of nucleases before use. (**B**) Urea-PAGE gels stained with SYBR show the effect of DNase I on samples collected during T7 Pol transcription assays and (**C**) assays that have FUS (2 μM) present. Gels shown are untreated samples (left) and those following incubation with DNase I (right). Control samples were incubated for 120 minutes with NTPs (-N) or T7 Pol (-T) during the transcription assay. Lanes that include an RNA control (Tet456) are labeled by the amount in nanograms (ng) loaded to the gel. Lanes with 1.5 and 5 ng Tet456, RNA was included during the transcription assay. Lanes with 25 and 100 ng Tet456 show RNA loaded to the gel from stock. Asterisks (*) indicate samples without DNase treatment. (**D**) Urea-PAGE gel images for samples collected during T7 Pol transcription (left) and those after treatment by RNaseA (right). (**E**) Replicates not already shown of time course experiments for T7 Pol transcription without (left) or with (right) 2 µM FUS protein added in the reaction (see also **Figures 1A** and **1B**, **Supplemental Figures 1B and 1C**). (#) indicates samples omitted from quantitation due to nuclease activity seen. (–) indicates lanes with no sample loaded. Controls are labeled as in (**B**). (**F**) Urea-PAGE gel images of transcripts produced by T7 Pol in titrating concentrations of urea, KCl, or NaCl. (**G**) Urea-PAGE analysis with SYBR-staining of replicates for transcription assays with increasing concentration of FUS (replicate 4 shown in Figure 1C). (**H**) Testing the role of protein aggregation in shifting RNA to the well during PAGE analysis. Samples were treated with either proteinase K, Prot K, or sodium dodecyl sulfate, SDS. (**I**) Replicates of Co-IP of FUS with T7 Pol. Replicates #1 is shown in part in **Figure 1D**. Lanes loaded with MBP-FUS and T7 Pol protein from stock are labeled “C”. Lanes labeled “F” indicates loading MBP-FUS only. “M” indicates molecular weight ladder.

**Supplemental Figure 2.**
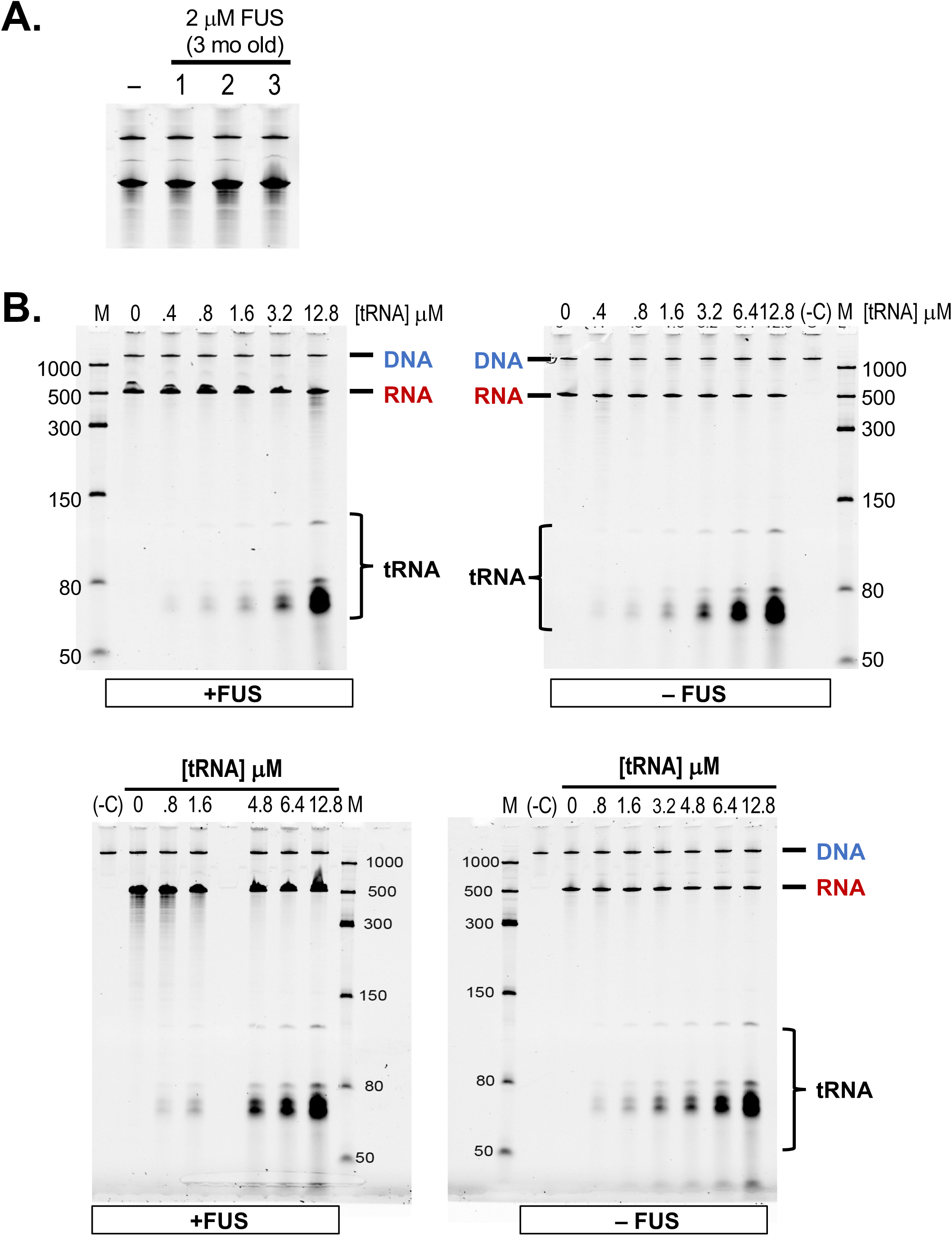
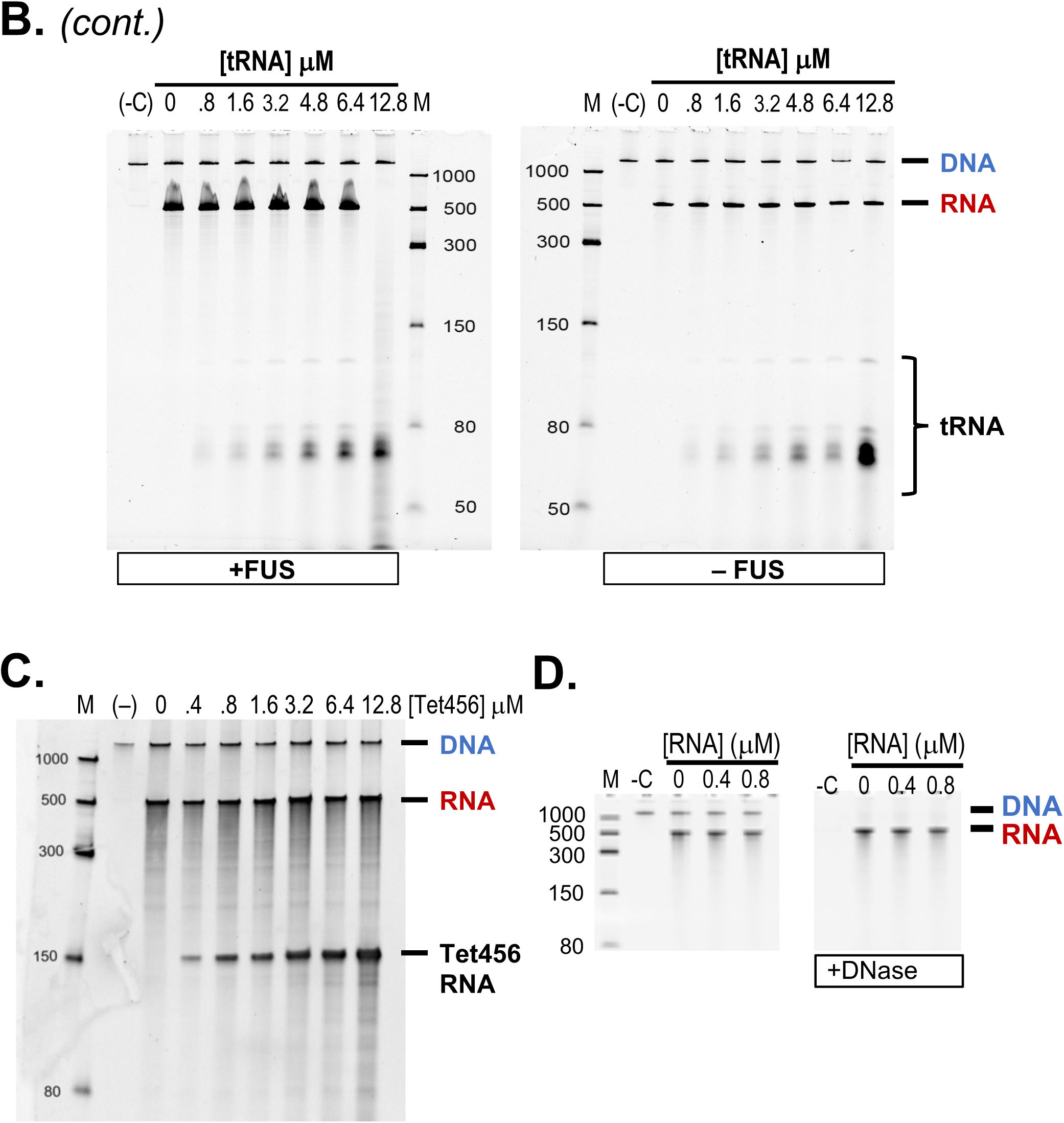
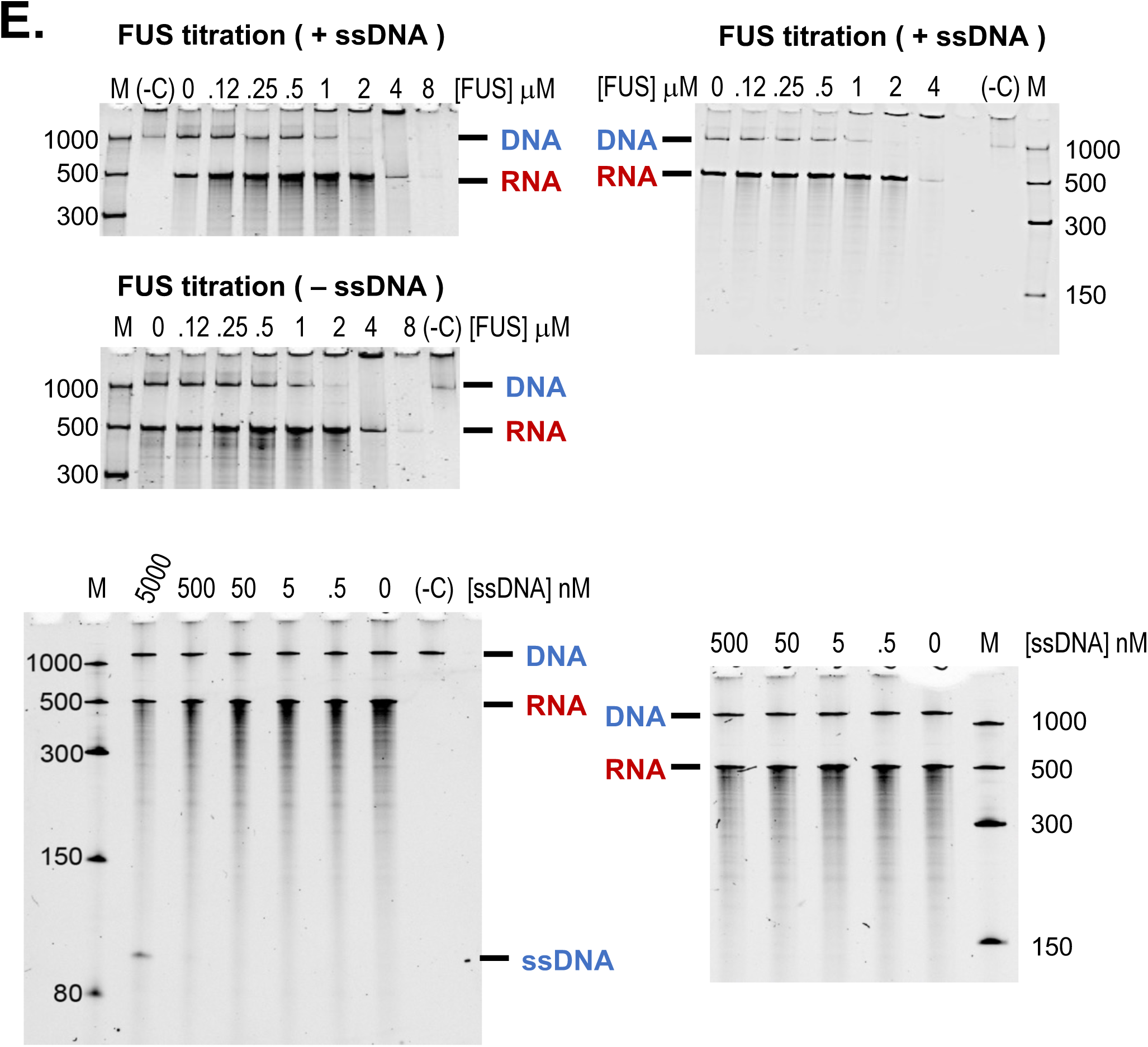
FUS activity is not competed by RNA or DNA. (**A**) Assay of the same FUS preparations used for **Figure 1B and 1C**, re-assayed after 3 months of storage at room temperature. (**B**) Titration of yeast tRNA into transcription run-off assays did not change the activity of FUS protein. Three replicate experiments are shown with 4 μM FUS (left) or no FUS (right) included in the reaction. (**C**) Titration of a TET456 RNA into transcription run-off assays did not change the activity of FUS protein. (**D**) Gel showing DNase treatment of samples with Tet456 RNA added. (**E**) Addition of single-stranded DNA, ssDNA, into transcription run-off assays did not change the titrated activity of FUS protein. Also shown are control experiments titrating ssDNA into T7 Pol transcription assays.

**Supplemental Figure 3.**
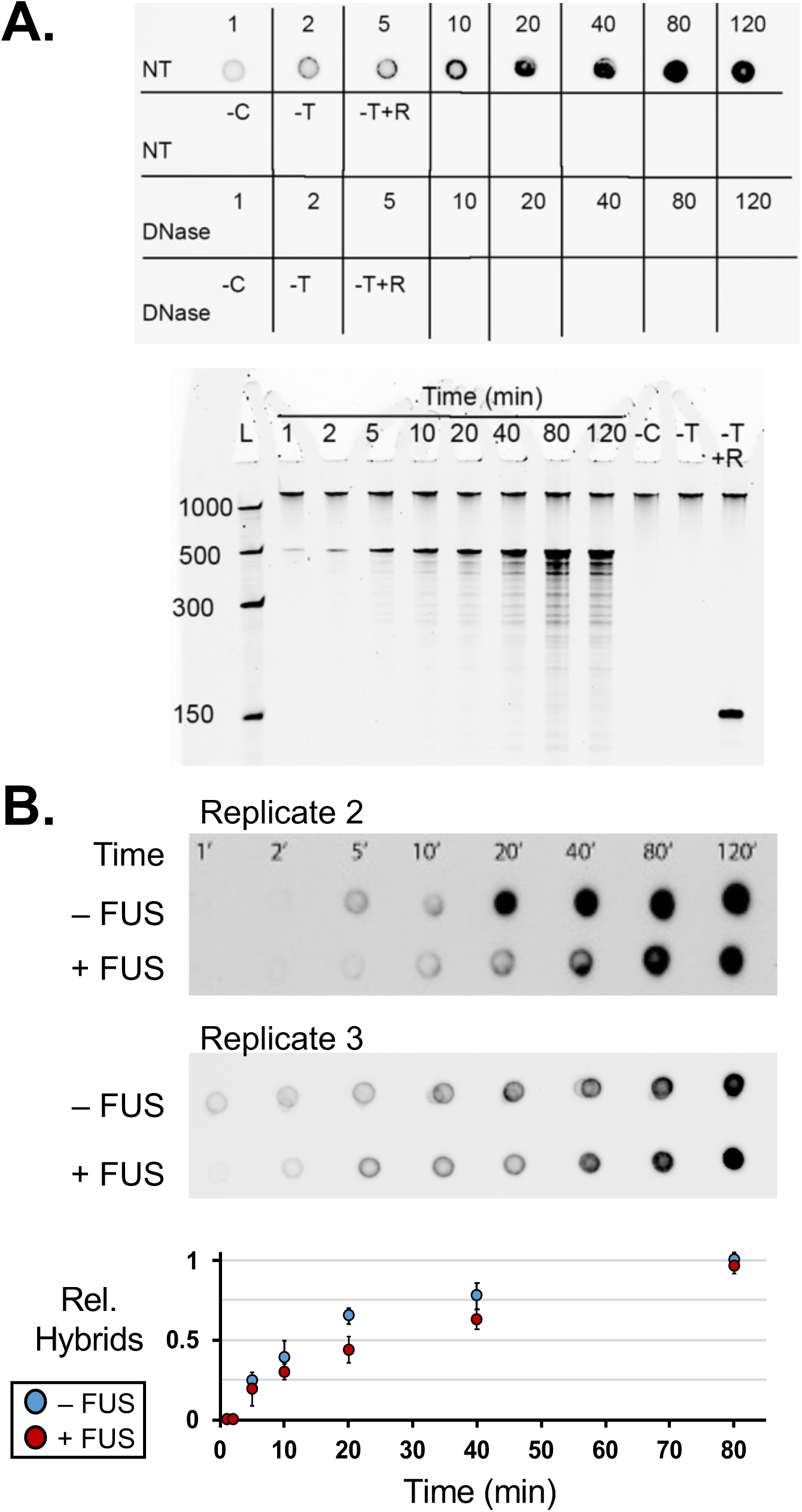
Time course of RNA:DNA hybrid formation during run-off transcription assays. (**A**) Assay as shown in **Figure 3A** and showing controls of omitting NTP, (–), or T7 Pol, -Pol, or addition of a non-specific RNA, +R. Samples were incubated with Dnase I to remove DNA and show the S9.6 antibody requires both RNA and DNA strands to bind. Note: enhanced Dnase I activity toward RNA:DNA hybrids was achieved by increased concentrations of MgCl_2_ (8 mM) and CaCl_2_ (1.3 mM). RNA products were detected by PAGE analysis and Sybr staining. The additional bad in the leftmost lane, -Pol+R, is that of TET456 RNA. (**B**) Replicates of dot blot assay shown in **Figure 3A** showing hybrids formed during T7 Pol run-off assays in the absence or presence of FUS (2 μM). Quantification of dot blot assays show hybrid levels differed most at timepoints collected at 20 and 40 min. Results are averaged from 3 experiments. Error bars represent standard error about the mean (±SEM).

**Supplemental Figure 4.**
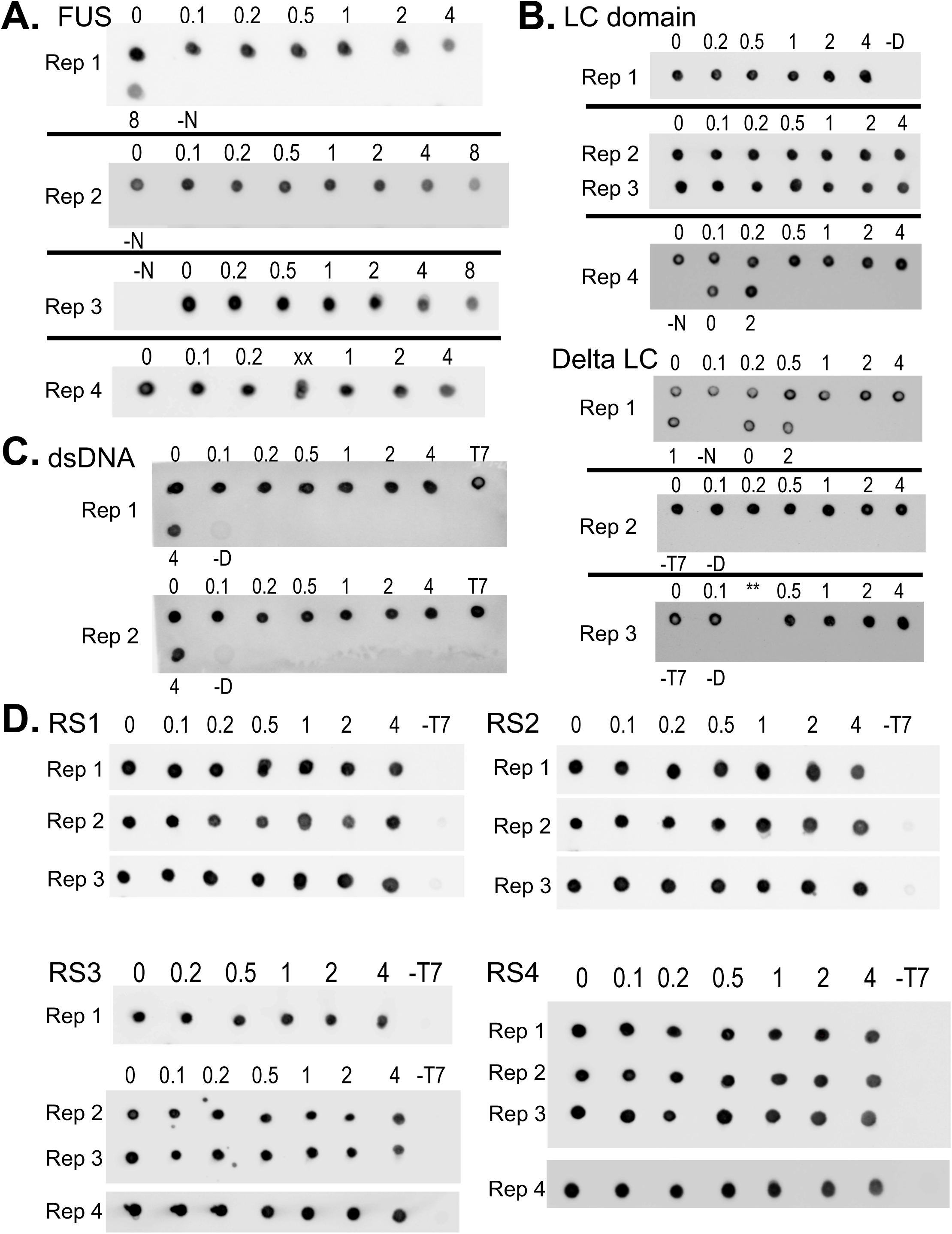
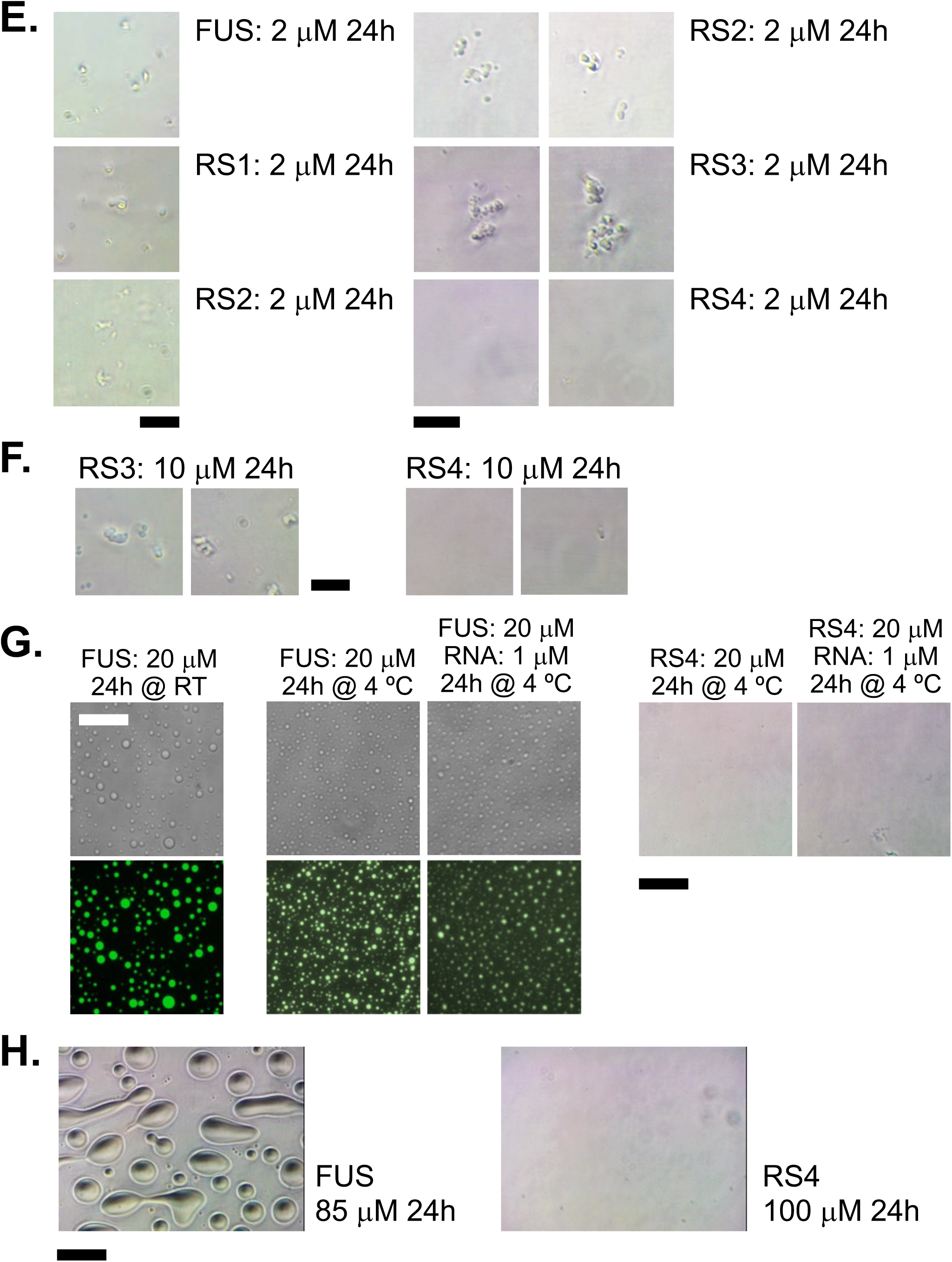
Replicate dot blot assays for RNA:DNA hybrid formed during run-off transcription assays and inspection of FUS particles using phase contrast microscopy. (**A**) Replicates of assays with titrating concentrations of FUS. Replicate 3 is included in **Figure 3B**. (**B**) Dot blot assays for FUS-LC and Delta LC titrations. (**C**) Dot blot assays for FUS-LC titrations using an antibody to double-stranded DNA (dsDNA). (**D**) Dot blot assays for arginine substituted FUS mutants RS1, RS2, RS3, and RS4. (**E**) Particles assembled after 24-hour incubation of FUS or RS proteins (2 µM) in transcription run-off assay conditions. (**F**) Imaging for particles formed by RS3 and RS4 proteins at 10 µM, showing no clear evidence of RS4 assembly. (**G**) Imaging for particles formed by FUS and RS4 proteins (20 µM) after incubation for 24 hours at 4 °C. Incubations of FUS and RS4 was repeated with addition of TET456 RNA (0.5 µM). Fluorescent images were made by addition of GFP-tagged FUS LC-domain (amino acids 1 to 266) at x150 lower concentration that FUS. (**H**) Imaging for particles formed by FUS or RS4 at high protein concentration as indicated. Scale bars are 2 µm for (**E**) and (**F**) and 4 µm in parts (**G**) and (**H**).

**Supplemental Figure 5.**
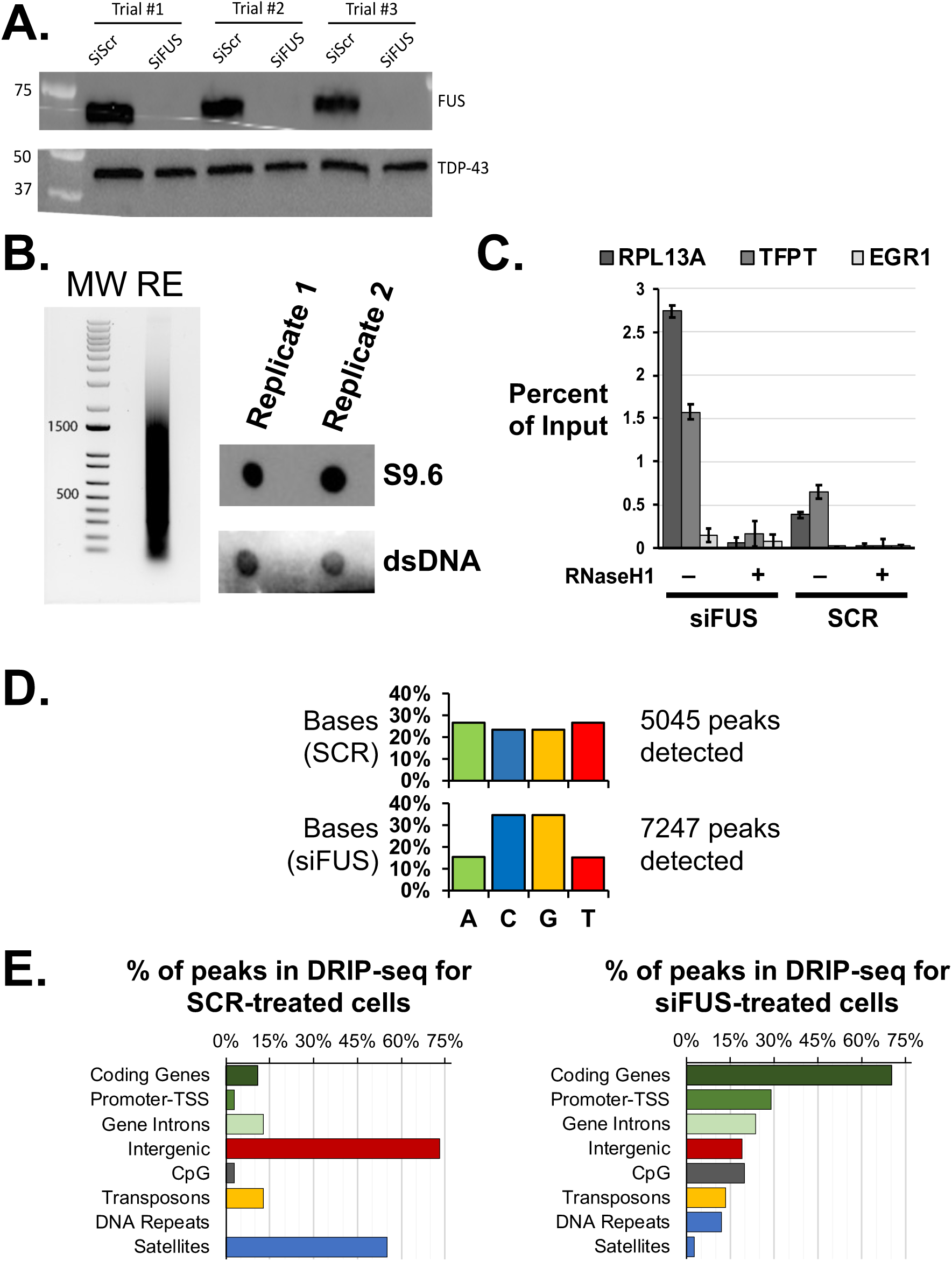
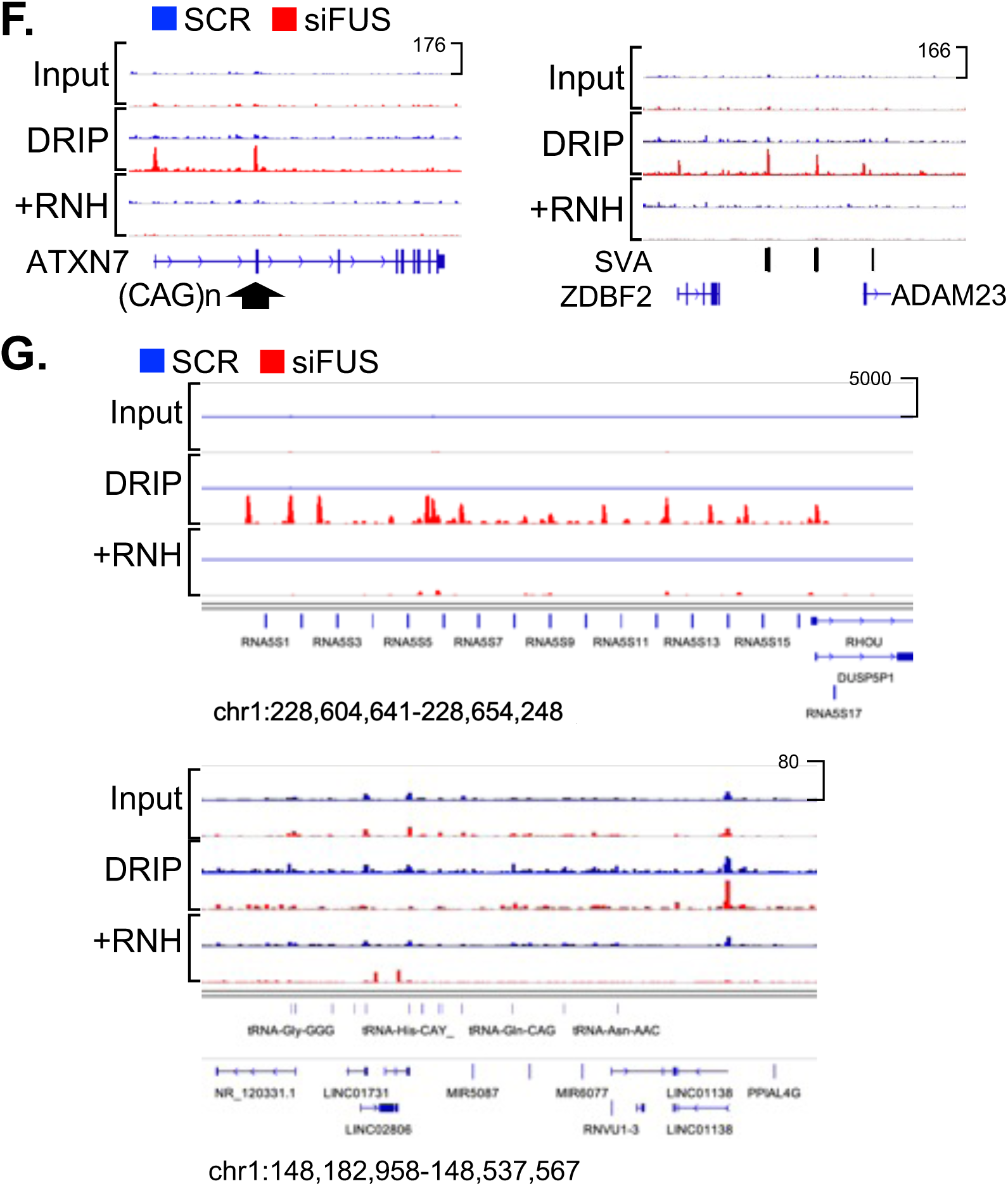
DRIP-seq analysis of FUS knockdown in HEK293T/17 cells. (**A**) Western assay to confirm knockdown of FUS protein by siFUS relative to control, SCR, in HEK293T/17 cells, with TDP-43 protein serving as loading control. See also **Figure 5.** (**B**) Verification of restriction enzyme digestion of genomic DNA recovered from HEK293T/17 cells, left, and the presence of RNA:DNA hybrids in genomic DNA recovered by dot blot assay, right, using the S9.6 antibody with antibody to dsDNA as loading control. (**C**) Realtime PCR to detect RNA:DNA hybrids after immunoprecipitation for R-loop positive promoters RPL13A and TFPT, and R-loop negative promoter EGR1. Samples were also treated with Rnase H, +RNH, prior to pulldown as negative control. Results shown are from 2 replicates and error bars indicate standard error about the mean (±SEM). (**D**) The base composition of DNA sequences in peaks called by MACS2 for DRIP-seq of SCR or siFUS treated cells. A GC content of >60% was found in peaks called for siFUS-treated samples. (**E**) Protein-coding genes comprised the largest percentage of peaks detected in the FUS knockdown results, in contrast to SCR-treated sample results. (**F**) DRIP-seq analysis of siFUS-treated samples reveals a peak in R-loop signal arising at a CAG-repeat DNA sequence at the *ATXN7* gene and SVA sequences, a SINE retrotransposon sub-type. (**G**) DRIP-seq signals observed at a 5S rRNA gene cluster and a tRNA gene cluster, both located on chromosome 1.

## Notes

https://www.ncbi.nlm.nih.gov/geo/query/acc.cgi?acc=GSE206740

### Competing Interest Statement

The authors have declared no competing interest.

### Summary of Updates

The text describing and discussing results has been rewritten for clarity. Additional data has been included, such as validation assays and replicates of figures shown in the main body.

